# Potential antiviral and immunomodulatory Amazonian medicinal plant compounds

**DOI:** 10.1101/2024.11.27.625749

**Authors:** Sarah Maria da Silva Napoleão, João Paulo Romualdo Alarcão Bernardes, Bernardo Guerra Tenório, Calisto Moreno Cardenas, Bruno Stéfano Lima Dallago, Serhat Sezai Çiçek, Roberto Messias Bezerra, Jorge Federico Orellana Segovia, Elida Cleyse Gomes da Mata Kanzaki, Isamu Kanzaki

**Author notes:** Correspondence should be addressed to: Colina Bloco H, AP 406. Campus Universitário Darcy Ribeiro, Universidade de Brasília, Asa Norte, DF. CEP 70.904-108. Brazil.

## Abstract

**Background:** Novel antiretroviral drugs are in constant need for HIV/AIDS patients to face the continuous emerging resistance to the commonly prescribed combination of anti-HIV synthetic agents and their side effects.

**Methods:** Amazonian medicinal plants, *Licania macrophylla* (Chrysobalanaceae) and *Ouratea hexasperma* (Ochnaceae) were assayed for antiretroviral and immunomodulatory activity, utilizing an established human leukocyte cell line and the Simian Immunodeficiency Virus. The IL-4, IL-6, IL-8, IL-10 and IFN-y cytokines were quantified after leukocyte culture stimulation with ethanolic plant extracts and challenged with lentivirus infection.

**Results:** Mitotic activity induced by *O. hexasperma* extract was significatively more pronounced than *L. macrophylla* extract. Cytokine modulation was recorded in SIV infected cells under independent treatment with *O. hexasperma* and *L. macrophylla* extracts. Betulinic acid, ninuriflavone, (-)epigallocathechine, (-)gallocatechine and 4-O-methyl-epigallocatechin were isolated from *L. macrophyla*.

**Conclusions:** Cellular proliferative activity and cytokine modulation by the extracts assayed have potential applications in the therapy of HIV/AIDS pathology, as the isolated compounds of these plants have been reported for antiviral activity.

## INTRODUCTION

Amazonian medicinal plants are a major source of therapeutic agents and active pharmacological compounds. Despite bioinformatic approaches applied to modify natural compounds and/or to design and synthetize new drugs, the biodiversity is still the most powerful and potential option to reveal new chemical compounds for a large array of prophylactic, therapeutic and cosmetic use. Anyhow, both strategies are applied for common purposes to rationally explore the natural resources utilization [1,2]. Infectious diseases, particularly of viral etiology, have their agents prone for vaccines development, however the high genetic variability of RNA viruses makes difficult the protective role of immunogens under investigation as for Dengue virus, HIV and SARS-CoV-2, among others, of epidemiological significance [3]. Chemical compounds that can interact with virus receptors or as ligands in cell receptors, display varied activities, therefore possible mechanisms in virus, glycoproteins and/or ligands, blocking its coupling to cell host receptors or else, as metabolites inducing the expression of cytokines are investigated [4].

The arsenal of antiretroviral drugs needs to be continuously implemented, mainly due to drug resistance and side effects development, commonly found among HIV infected subjects and related diseases as AIDS, under antiretroviral therapy [5,6]. Host defense system to virus infection initiates with the organisms physical, chemical and biological barriers for the invasive agent, and can continue with the innate immune response progressing or not to the adaptive immune response. Biological processes of immune response usually proceed with cytokine signaling to activate the effector mechanisms of both innate and adaptive immune response [7,8]. Cytokines mediate the inflammatory and anti-inflammatory host response during the course of infection and usually, for healthy subjects, the morbidity is resolved [9]. Nowadays, as a common scenario, mainly in urban areas, people are deprived of a health environment as also adequate nutrition, mental and emotional balance [10]. Human depression is prevalent in large metropolis, all around the world, which is immunologically characterized by the secretion of pro-inflammatory cytokines as IL-1ß, TNF-α and IL-6 [11] besides, chronic inflammation predisposes to a large array of diseases, mainly linked to poor immune response to infectious agents [12]. Amazonian biodiversity provides a bunch of resources for human health as natural products, among them, medicinal plants that are commonly used by traditional populations, for the treatment of their maladies. Folklore and knowledge of ancient civilizations are hidden concerning the scientific data of plants used, just remaining the procedures the plants are prepared and for which ailment. In order to rescue chemical, biological and medical information of these plants, systematic listing of used plants, their geographical distribution and preliminary chemical and biological studies have been conducted [13–17].

*Licania macrophylla* (Benth), meanwhile reclassified as *Hymenopus macrophyllus* (Benth), by Sothers & Prance [18] in the Chrysobalanaceae botanical family, is popularly known as “anauerá”, derived from a native amerind language in the Amazon. *L. macrophylla* exhibits a typical bush morphology, of 17-28 meters height, growing in the margin of inundated areas. Besides its medicinal use for the treatment of gastrointestinal disorders of infectious etiology, the timber is employed for construction purposes [19]. Anauerá chemical compounds have been isolated and characterized, as the licanol, a flavanol: (-)-4’-O-methyl-epigallocatechin-3’-O-α-L-rhamnoside [20]. Previously, we reported the inhibitory activity of *L. macrophylla* ethanolic extract (bark) for multidrug resistant *Staphylococcus aureus* and *Pseudomonas aeruginosa* [21].

Commonly known as “barbatimão do cerrado” among natives in northern Brazil, it is taxonomically classified as *Ouratea hexasperma* (A. St. -Hil.) Baill in the Ochnaceae botanical family. Its shrubby structure reaches 5 meters height with wrinkled bark and tortuous branches, typical of “cerrado” vegetation (Brazilian savannah) [22]. Phytochemical analysis carried out by Daniel et al. [23] revealed the presence of 7”-O-methylagathisflavone, agathisflavone, epicatechin and the mixture of 6-C-glycopyranosyl-luteolin and 3-O-glycopyranosyl-quercetin. Also, Fidelis et al. [24] reported the isolation of trans-3-O-methyl-resveratrol-2-C-β-glucoside, lithospermoside, 2,5-dimethoxy-p-benzoquinone, lup-20(30)-ene-3β,28-diol, 7-O-methylgenistein, apigenin and luteolin and amentoflavone. The above characterized compounds and crude extracts of *O. hexasperma* exhibited in vitro varied biological activities as antioxidant, anticarcinogenic, antidiabetic, antimicrobial, anti-inflammatory and so on [17,24,25].

## MATERIALS AND METHODS

### Plant extracts

Botanic samples of *Licania macrophylla* (Chrysobalanaceae) and *Ouratea hexasperma* (Ochnaceae) were collected in low lands surrounding the Amapari river, at the geographic coordinates 0° 46’ 14” N/51° 55’ 54” W (Pedra Branca do Amapari municipality), and in the Amapa “cerrado” (savannah) at the geographical coordinates 0°55’51” N/51°11’35”W (Ferreira Gomes municipality), respectively (Fig. 1). Hundred grams of each, dried and pulverized, plant material, leaves of *Licania macrophylla*, and bark of *Ouratea hexasperma*, were macerated in 1 L of ethanol for 7 days at room temperature, repeated for three times the procedure. The extract was obtained in a rotary evaporator (Quimis) at 40 °C under low pressure, and kept under refrigeration until use. Plant materials of *Ouratea hexasperma* and *Licania macrophylla* were deposited at the Institute of Scientific and Technological Research of Amapa State under Registration number 16593 and 16594, respectively.

**Figure 1.**
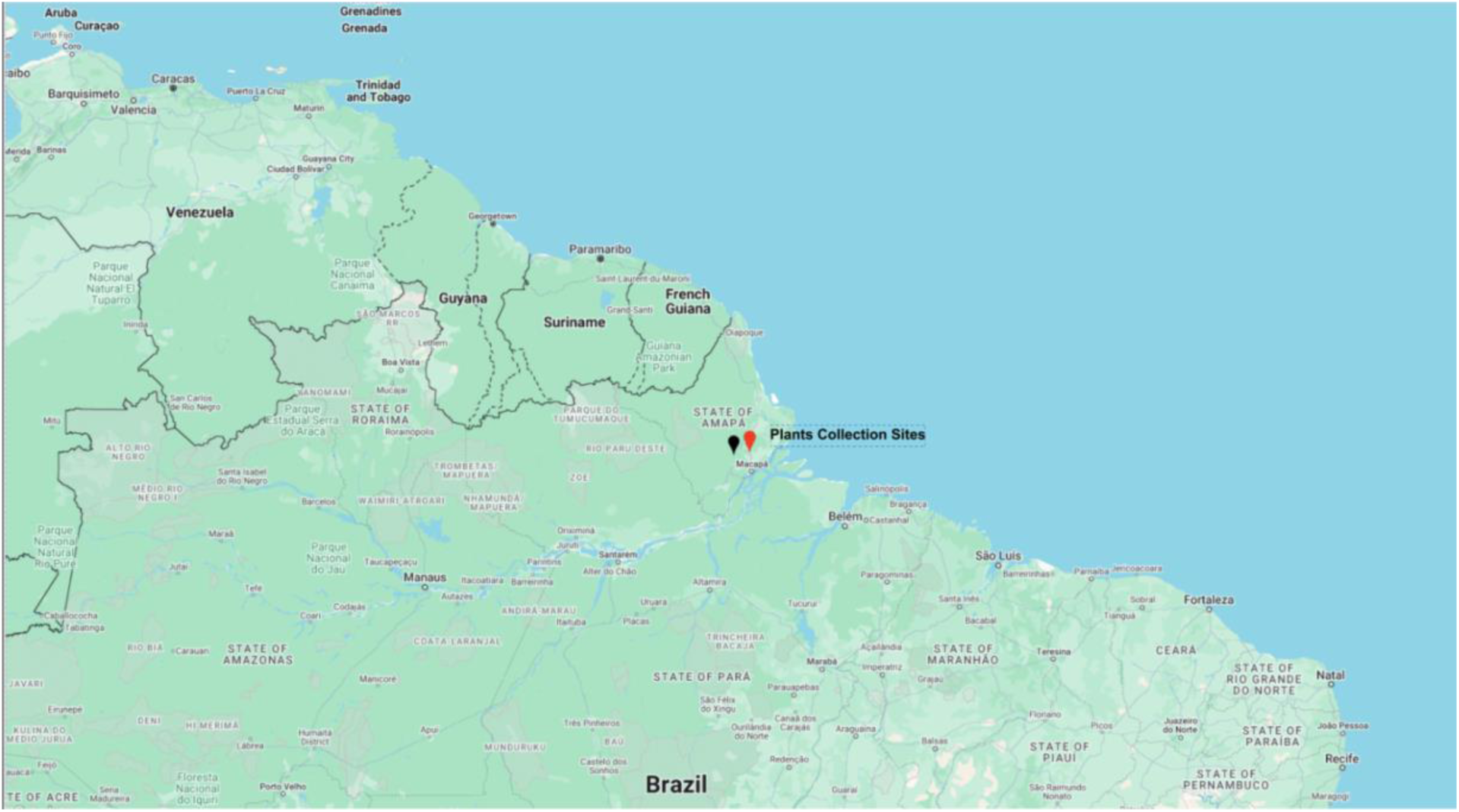
Sites of plant material collection in the Amapa state, northern Brazil. Black pin: Pedra Branca do Amapari municipality; red pin: Ferreira Gomes municipality.

### Ultra-High-Performance Liquid Chromatography (UHPLC) Analysis

The methanolic extract of *Licania macrophylla* was analyzed by ultra-high-performance liquid chromatography (Shimadzu Nexera) composed of a controller (CBM 20-A), a degasser (DGU-20A), two binary pumps (Nexera X2 LC-30AD), an autoinjector (Nexera X2 SIL-30AC), a thermostable column compartment (CTO-20AC), a PDA/UV detector (SPD-M20A), and a triple quadrupole mass spectrometer (LC-MS-8030) equipped with an electrospray ionization (ESI) source. Separation was achieved on a C18 column of 100 x 2.1 mm and 1.6 μm particle size (Luna Omega Polar, Phenomenex). The data were processed using LabSolutions LCMS software version 5.96 (Shimadzu Corporation) [13,15].

### Cell viability assay

Human leukocytes transformation was carried out as described elsewhere [26,27]. Briefly, leukocytes were obtained from total heparinized blood by centrifugation in sucrose gradient (Histopaque-1077, Sigma), maintained in RPMI medium supplemented with 10 % fetal bovine serum, 1 % antibiotics (penicillin and streptomycin, Sigma) and transformed by the addition of 10% of Hut-78 [28] cell culture supernatant and cultivated in an incubator (Sanyo) at 37°C, 5% CO_2_ atmosphere and humidity. Established human leukocyte culture (EHLC) was treated with multiple serial dilutions of each extract. Cell viability was assessed by trypan blue staining (Sigma), showing and counting damaged/dead cells, as also by spectrophotometric measurement, in cell culture supernatant, of the mitochondrial succinate tetrazolium reductase enzyme in a commercial cytotoxic assay kit (CytoSkan WST-1, Roche).

### Antiretroviral assay and cytokine quantification

EHLC, cultivated in 25 cm^2^ flasks with 5X10^5^ cells/mL, was treated with low cytotoxic plant extract concentration as above determined. After 24 hs plant extract treatment, cells were infected with 100 μL of SIVmac_251_ (NIH AIDS Research Reagents, US) at 599.64 pg/mL of SIVp27 concentration. Cell supernatant samples were harvested at post viral infection intervals of 48 hs, 144 hs and 240 hs, cleared by centrifugation and kept at -20 °C until use for antiretroviral activity by quantification of SIVp27 antigen (SIVp27 Antigen Capture Assay, Advanced Bioscience Laboratory, Inc), as also for IL-4, IL-6, IL-8, IL-10 and IFN-y quantification (DuoSet ELISA Development System). Equal amount of fresh medium, 3 mL, was added after each cell supernatant harvesting. SIVmac_251_ was previously produced in Hut-78 cells and viral antigen (SIVp27) purified from cells’ supernatant. For determination of viral antigen concentration, briefly, 25 μL of disruption buffer was added to each microtiter well coated with monoclonal antibody to SIVp27 antigen followed by the addition of 100 μL of cell’s supernatant to each well, as also a serial dilution of SIVp27 standard antigen (2,000-62.5 pg/mL), including the negative control consisting of complete RPMI medium. After 60 min. incubation at 37 °C, wells were washed and 100 μL of conjugate solution (horseradish peroxidase-labeled, mouse monoclonal antibody to SIVp27) was added to each well and incubated again. After 60 min., wells were washed, and added 100 μL of peroxidase substrate. After 30 min. incubation at room temperature, the chromogenic reaction was blocked by the addition of 2N sulfuric acid solution. Plates were read at 450 nm in a spectrophotometer, microelisa plate reader (Thermoplate) and absorbance registered as optical density.

### Cytokine quantification

EHLC treated and cultivated with optimum plant extract concentration (low toxicity), and infected after 24 hs plants’ extract treatment, with SIVmac251, had supernatant samples harvested at intervals of 48 hs, 144 hs and 240 hs. The assay controls were supernatant of SIV uninfected and plant extract untreated cells, SIV uninfected and plant extract treated cells. Plate wells were sensibilized overnight at 4 °C with capture antibody to each interleukin (IL-4, IL-6, IL-8, IL-10 and IFN-y), diluted in PBS according to the kit instructions (Duo Set-R&D-Systems, USA). After 4 times washing of wells with buffer, the blocking solution (PBS-BSA 1%-0,05% NaN_3_) was added, and after 1 hour incubation at room temperature, the wells were washed with buffer, and reference interleukins and leukocyte cell culture supernatant were added to each well, in triplicate, and incubated at room temperature for 2 hours. Diluted detection antibody solution was added to each previously washed well, and at the end of 2 hours incubation at room temperature, streptavidin-peroxidase solution was put to each well and washed after 20 min. incubation for the addition of the chromogenic substrate solution (TMB), protected from light exposure, and incubated for 30 min. Sequentially, the reaction was stopped with sulfuric acid solution and the absorbance registered at 450 nm in a multi-plate spectrophotometer reader, expressed in optical density (Thermoplate). These results were confirmed by immunoassays for the same cytokines above analyzed, utilizing ELISA kits kindly donated by the ImmunoTools GmbH company.

### Statistical analysis

Obtained data was analyzed utilizing the Graphpad prism software 10.0 version submitted to Two Way ANOVA and appropriately applying the Tukey’s multiple comparison test.

## RESULTS

The UHPLC-PDA-MS analysis revealed betulinic acid and niruflavone as the most abundant compounds in the leaf extract of *L. macrophylla*. Additional compounds identified by comparisons with the literature data were (-)-gallacatechin, (-)-epigallocatechin, and (-) -4’-O-methyl-epigallocaterchin.

Previously, we did not detect cytokine production in Hut-78 cell line supernatant after stimulation with plant extracts and SIV infection, so the EHLC was generated to carry out the experiments to detect cytokine production.

The optimal plant extract dilution was determined by the combined results of chemical (cytotoxic assay) and phenotypic (cell counting/trypan blue staining) analysis. An optimal dilution of 1:1024, for both botanical species corresponding to 0.34 μg/mL, was utilized to treat the EHLC in culture. Cells were counted during time intervals from 48 hs to 288 hs, post plants’ extracts treatment. Concerning *L. macrophylla* extract treatment, the initial and final time intervals of 48 hs and 288 hs, treated cells had 1.5 times more cells than the control (1.5X10^5^ treated cells versus 1.0X10^5^ untreated cells), and 2.3 times more cells than the control (1.6X10^5^ treated cells versus 0.7X10^5^ untreated cells), respectively, with statistical significative differences (p=0.0133 and p=0.0122, respectively). In the time interval of 96 hs, there is not a significative difference in cell density between treated and untreated cells (p=0.0842). In the intermediate intervals of 144 hs, 192 hs and 240 hs, *L. macrophylla* extract treated cells grew less than the control with statistical significative differences (p=0.0069, p=0.0177 and p=0.0039, respectively). Cells treated with *O. hexasperma* extract had exceptional cell growth population after 48 hs and 96 hs post treatment, comparing to the control, about 3.3 times and 1.4 times higher (p=0.0182 and p=0.0087), respectively. In the time interval of 192 hs, treated cells growth dropped from 4.6X10^5^ to 2.8X10^5^ cells/mL, while the untreated cells kept at 4.4X10^5^ cells/mL (p=0.0087). In the final time interval of 288 hs, *O. hexasperma* extract treated cells kept growing, with 1.5X10^5^ cells/mL while untreated cell population growth dropped to 0.5X10^5^ cells/mL (p=0.0033). Comparing the mitotic activity of cells treated with the extracts of *L. macrophylla* and *O. hexasperma*, overall *O. hexasperma* extract treated cells showed higher cell density than *L. macrophylla* extract treated cells. From 48 hs to 192 hs time intervals, the mitotic activity difference between *O. hexasperma* extract and *L. macrophylla* extract reduced cell density from 2.4 times to 1.5 times, favorable to *O. hexasperma* extract treatment (p=0.018, p=0.0074 and p=0.0170, respectively). In the final intervals of 192 hs to 288 hs, there is a small cell proliferation difference (p=0.1001 and p=0.0125) between the treatments (Fig. 2).

**Figure 2.**
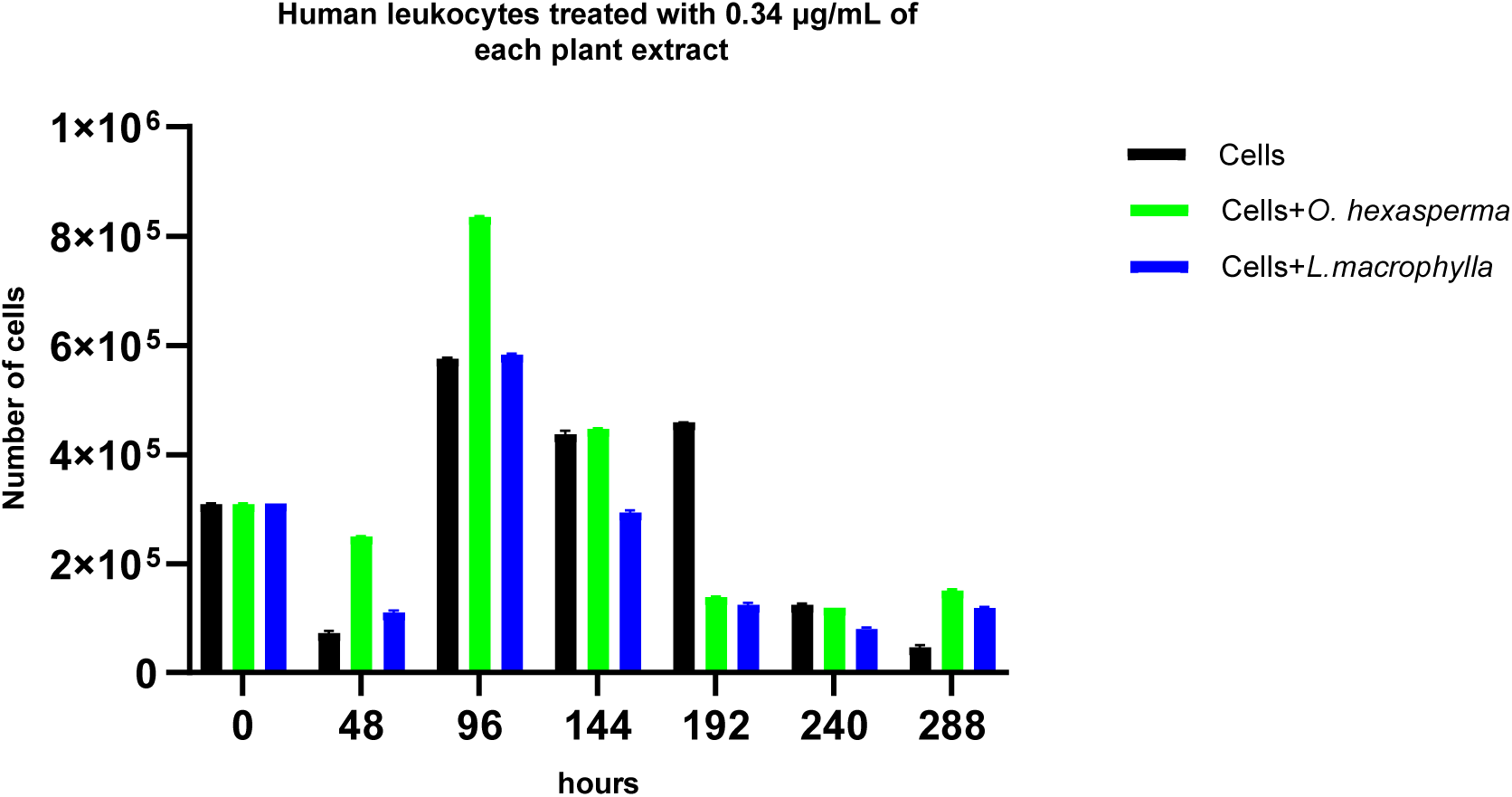
Established human leukocyte cell line population growth under treatment of *L. macrophylla* and *O. hexasperma* extracts evaluated during 288 hs culture. Initial time interval of 0 hs, all cells, untreated and treated had the same population density.

EHLC protection from lentivirus infection, SIVmac_251_, utilizing *L. macrophylla* and *O. hexasperma* extracts treatment, based on cell survival and cytopathic effects, showed that *L. macrophylla* extract treatment had significatively protective effects at 144 hs and 192 hs time intervals (p=0.0127 and p=0.0476, respectively), while *O. hexasperma* extract treatment reduced viral cytopathic effect on cells at 96 hs time interval (p=0.0144). In the final time intervals of 240 hs and 288 hs, both extracts treatment had no protective effect on the cells (Fig.3).

**Figure 3.**
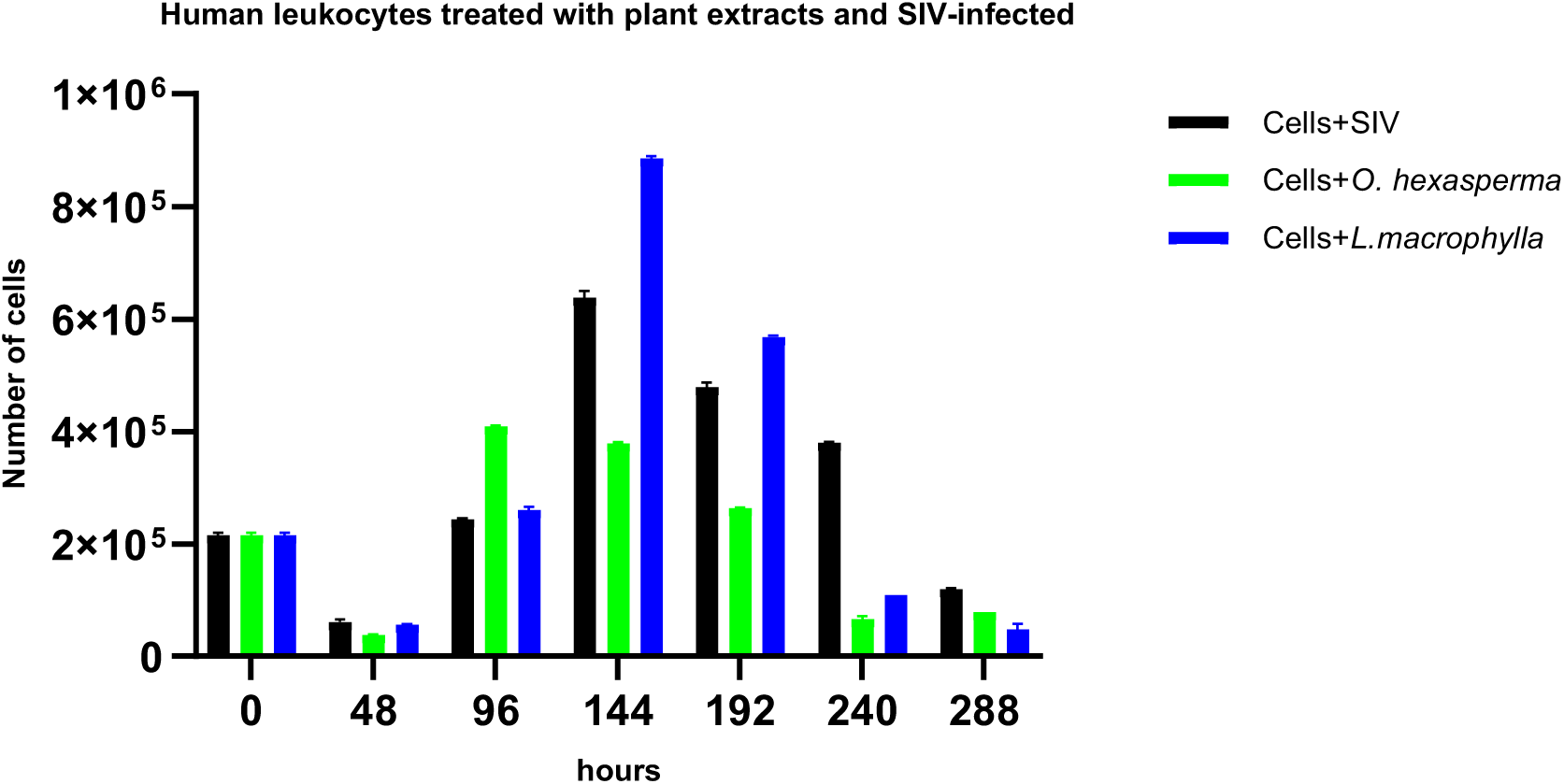
Established human leukocyte cell line treated for 24 hs, independently with *L. macrophylla* and *O. hexasperma* extracts and subsequently infected with SIVmac251. Cell viability was accessed until 288 hs of culture. The initial time interval of 0 hs, all cells had the same population density.

Cytokine expression by cells after treatment with plants extracts, and challenged with lentivirus infection, SIVmac_251_, showed the following patterns. Concerning cells culture under *O. hexasperma* extract treatment and SIV infection, IL-4 expression (Fig. 4) was significatively higher at 48 hs time interval in comparison to all other treatments (p ranging from 0.0208 to 0.0210), except in *L. macrophylla* extract treatment (p=0.1057). At 144 hs time interval, it was significatively (p ranging from 0.0005 to 0.0022) higher than the other treatments, except when comparing to *L. macrophylla* extract treatment and SIV infection, and the solely treatment with *L. macrophylla* extract (p ranging from 0.2736 to 0.3056). At 240 hs, it was significatively higher comparing to all treatments and SIV infection (p ranging from 0.0197 to 0.0205). Cells culture under treatment with *L. macrophylla* extract and SIV infection did not display any significative difference in IL-4 expression comparing to all other treatments, during all time intervals. In the initial stage of the experiment, differences were not significative for cells solely treated independently with both extracts and when infected by SIV (p>0.9999). In all other conditions, cells extract treated compared to cells extract treated and SIV infected, there were significative statistical differences (p=0.0040).

**Figure 4.**
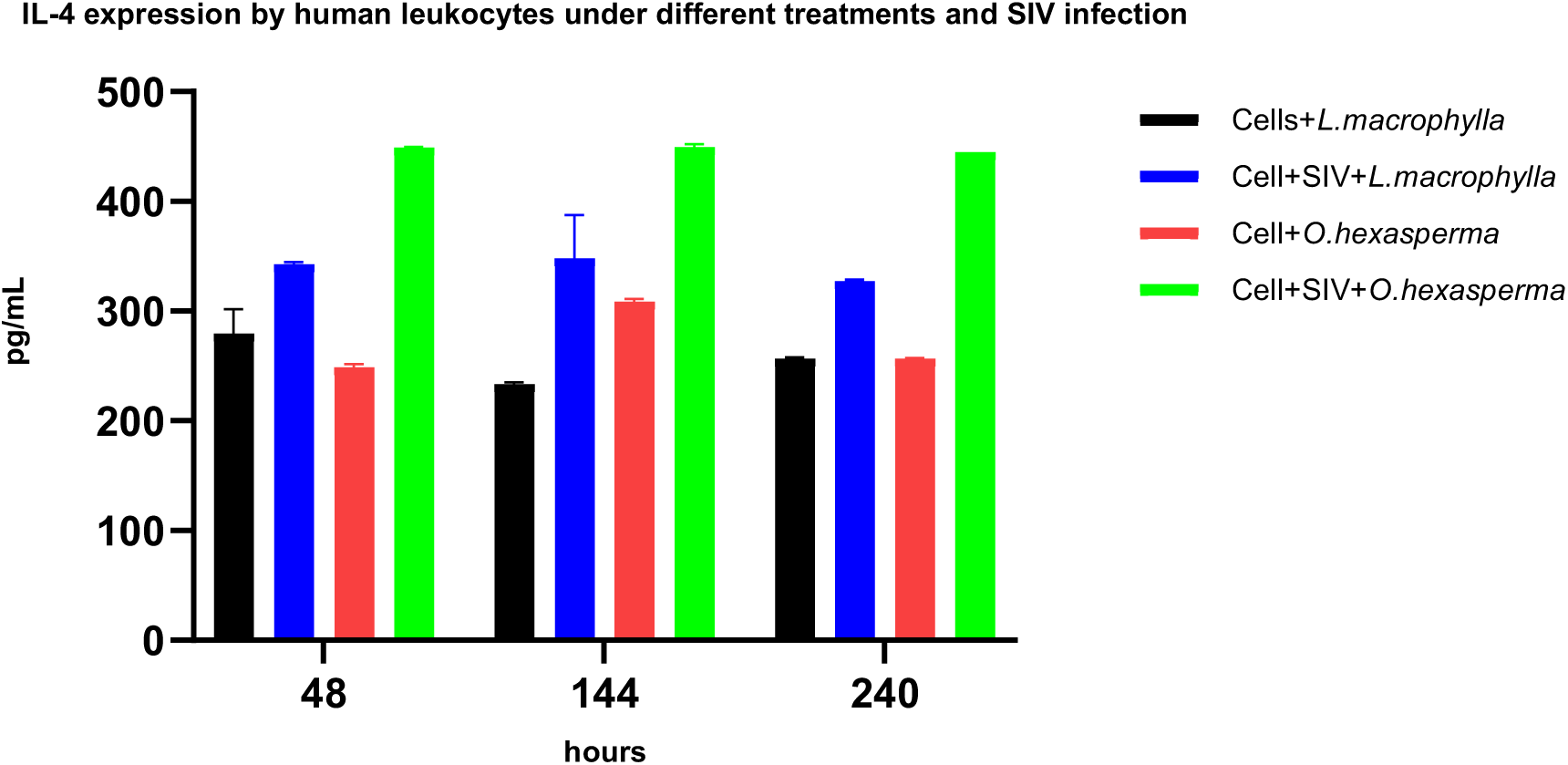
Interleukin 4 expression in established human leukocyte cell line treated with *L. macrophylla* and *O. hexasperma* and SIV infected, and in controls of cells submitted to extracts treatment and no SIV infection.

IL-6 expression (Fig.5) by the EHLC at 48 hs time interval yielded significative difference (p=0.0006) with higher IL-6 secretion by *L. macrophylla* extract treated cells and SIV infected than cells solely treated with *L. macrophylla* extract. Comparing both SIV infected cells but previously and independently treated with *L. macrophylla* and *O. hexasperma* extracts, showed that cells treated with *O. hexasperma* extract and SIV infected had significatively (p=0.0030) high IL-6 secretion. Cells solely treated with *L. macrophylla* significatively (p=0.0033) produced more IL-6 than cells treated with *O. hexasperma* extract and SIV infected. Cells solely treated with *L. macrophylla* extract had significatively (p=0.0049) higher IL-6 production than *O. hexasperma* extract treated cells. Other cell’s treatments with and without SIV infection (*O. hexaspema* extract treatment and SIV infection versus *O. hexasperma* extract treatment; *L. macrophylla* extract treatment and SIV infection versus *O. hexasperma* extract) did not show any statistical significative difference (p ranging from 0.0748 to 0.1467). In the following time interval of 144 hs, the highest significative difference (p<0.0001) in IL-6 secretion was achieved by cells solely treated with *O. hexasperma* extract compared to cells treated with *L. macrophylla* extract and SIV infected, as also when comparing to cells treated with *O. hexasperma* extract and SIV infected (p=0.011), and cells solely treated with *L. macrophylla* extract (p=0.0104). Cells solely treated with *L. macrophylla* extract had significatively higher production of IL-6 than cells treated with *O. hexasperma* extract and SIV infected (p=0.0095). The other treatments (*L. macrophylla* extract treatment and SIV infected versus *O. hexaspema* extract treatment and SIV infected; *L. macrophylla* extract treatment and SIV infected versus *L. macrophylla* extract treatment) did not result in any statistical significance (p ranging 0.0512 to 0.1718). At the last time interval of 240 hs, cells treated with *O. hexasperma* extract and SIV infected produced significatively more IL-6 than cells treated with *L. macrophylla* extract and SIV infected (p=0.0013) as also in cells solely treated with *O. hexasperma* extract (p=0.0177). Cells only treated with *L. macrophylla* extract had higher significative production of IL-6 than cells treated with *L. macrophylla* extract and SIV infected (p=0.0017), as also in cells only treated with *O. hexasperma* extract (p=0.0096). Comparison between cells only treated with *L. macrophylla* extract and cells treated with *O. hexasperma* extract and SIV infected did not show any significative statistical difference (p=0.0907).

**Figure 5.**
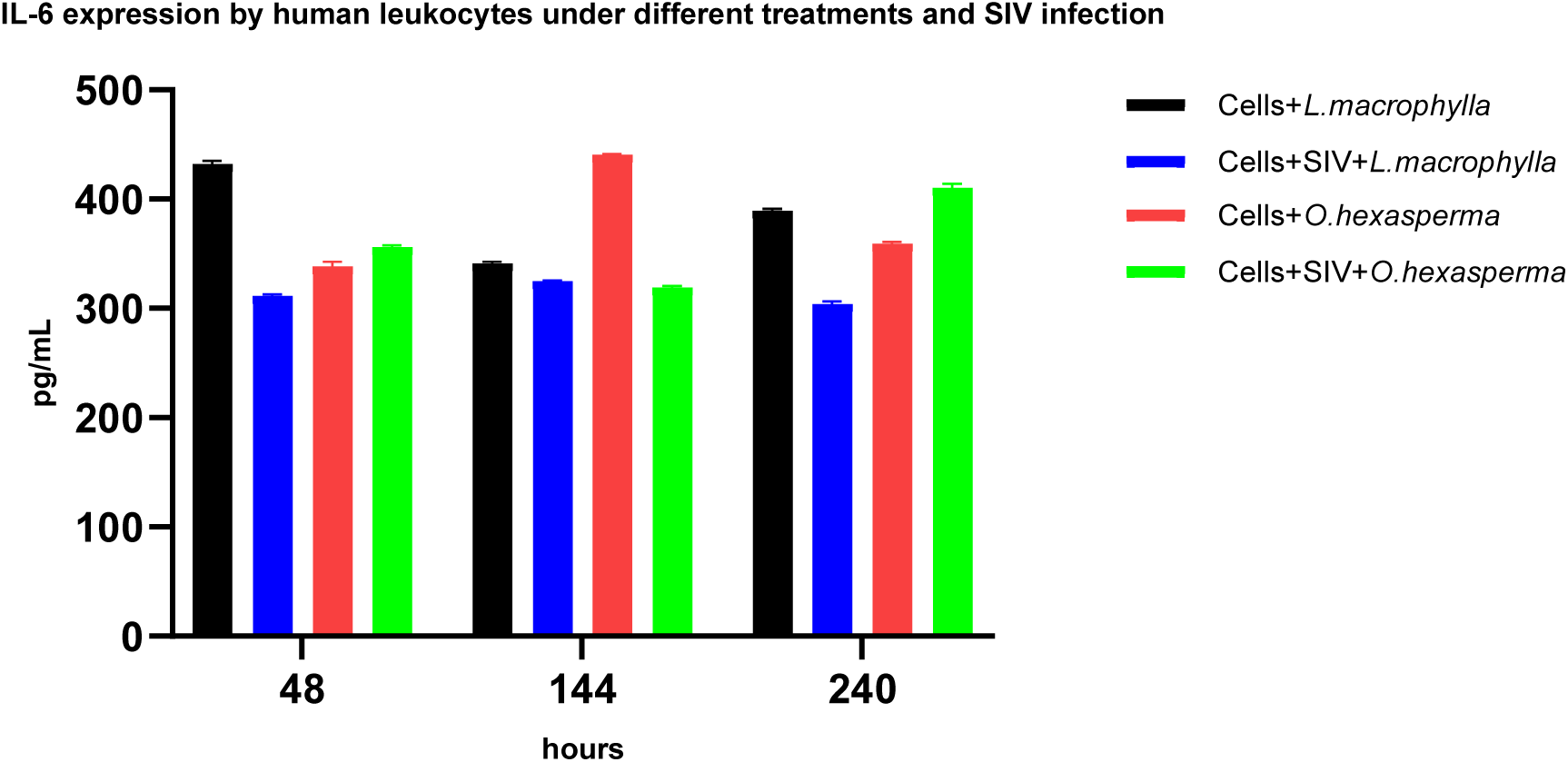
Interleukin 6 expression in established human leukocyte cell line treated with *L. macrophylla* and *O. hexasperma* and SIV infected, and in controls of cells submitted to extracts treatment and no SIV infection.

At 48 hs time interval, EHLC treated with *L. macrophylla* extract and SIV infected significatively produced more IL-8 (Fig. 6) than cells treated only with *O. hexasperma* extract (p=0.0241), as also in cells treated with *O. hexasperma* extract and SIV infected (p=0.0113). Cells solely treated with *L. macrophylla* extract secreted significatively more IL-8 than cells only treated with *O. hexasperma* extract (p=0.0248). The other treatments (*O. hexasperma* extract treatment versus *O. hexasperma* extract treatment and SIV infection; *L. macrophylla* extract treatment versus *O. hexasperma* extract treatment and SIV infection; *L. macrophylla* extract treatment versus *L. macrophylla* extract treatment and SIV infection) did not show any statistical significative difference (p ranging from 0.0530 to 0.1031). Sequentially, at 144 hs time interval, cells solely treated with *O. hexasperma* extract secreted more IL-8 than cells only treated with *L. macrophylla* extract (p=0.0058), as also in cells treated with *O. hexasperma* extract and SIV infected (p=0.0447). Cells treated with *L. macrophylla* extract and SIV infected significatively expressed more IL-8 than cells solely treated with *L. macrophylla* extract (p=0.0195) as also in cells treated with *O. hexasperma* extract and SIV infected (p=0.0386). Comparison of cells treated with *O. hexasperma* extract infected or not with SIV, and cells treated with *L. macrophylla* extract and infected or not with SIV, did not show any statistical significative difference (p ranging from 0.0607 to 0.1541). Lastly, at 240 hs, leukocytes treated with *O. hexasperma* extract and SIV infected, significatively secreted more IL-8 than cells solely treated with *O. hexasperma* extract (p=0.0036), as also in cells only treated with *L. macrophylla* extract (p=0.0016). The other conditions (*L. macrophylla* extract treatment versus *O. hexasperma* extract treatment; *L. macrophylla* extract treatment versus *L. macrophylla* extract treatment and SIV infection; *L. macrophylla* extract treatment and SIV infection versus *O. hexasperma* extract treatment and SIV infection) had no significative statistical difference in IL-8 expression (p ranging from 0.0525 to 0.2367).

**Figure 6.**
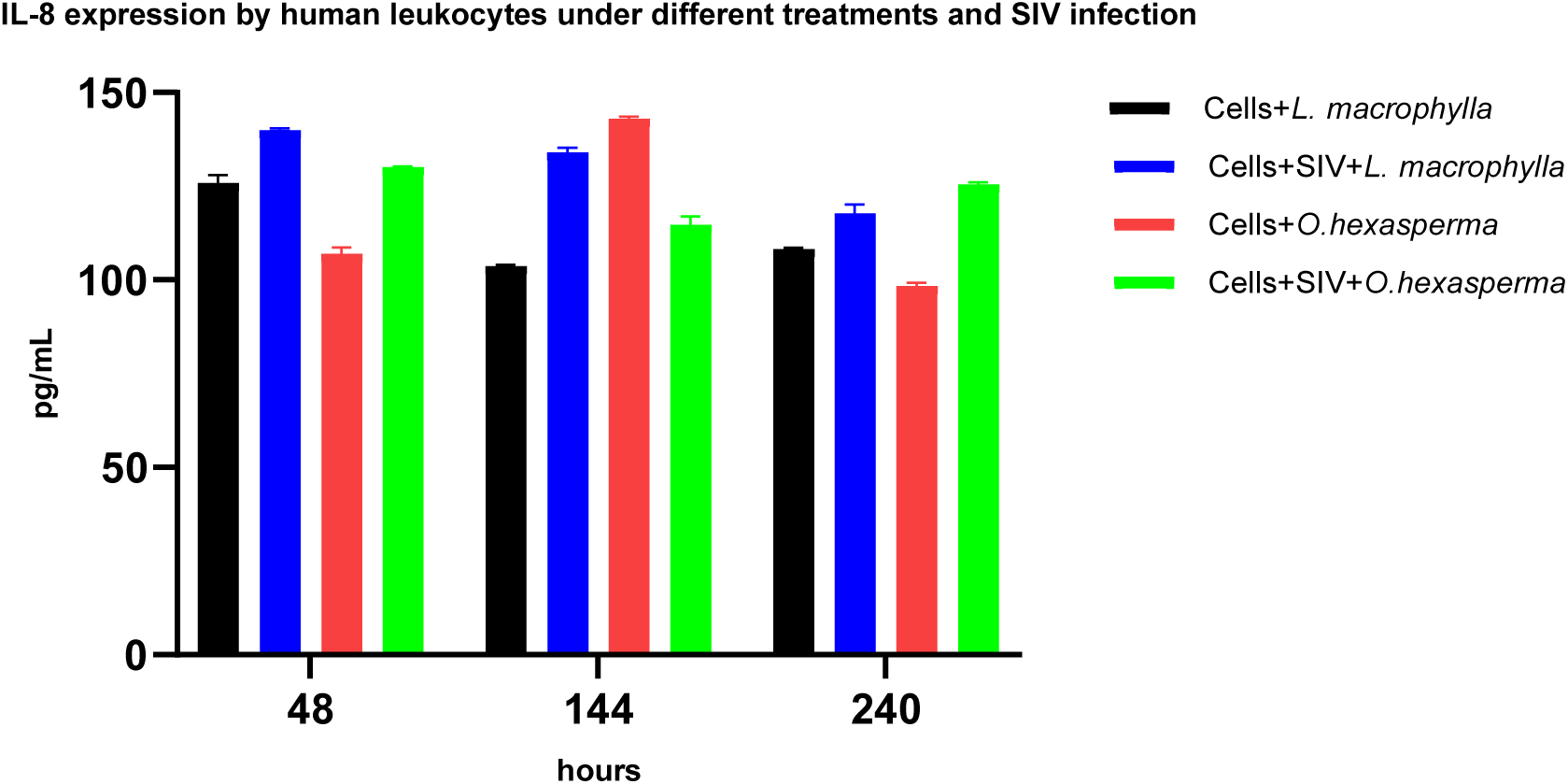
Interleukin 8 expression in established human leukocyte cell line treated with *L. macrophylla* and *O. hexasperma* and SIV infected, and in controls of cells submitted to extracts treatment and no SIV infection.

Just cells treated with *L. macrophylla* extract, at 48 hs, secreted IL-10 (Fig. 7), while in the other conditions the cytokine was not detected. At 144 hs time interval, cells treated solely with *L. macrophylla* extract significatively produced more IL-10 than cells treated with *O. hexasperma* extract (p=0.0156) and cells treated with *L. macrophylla* and SIV infected (p=0.0209), while in the other treatment the cytokine production was inexpressive. At the last time interval of 240 hs, cells treated with *L. macrophylla* extract remained producing more IL-10 than the other treatments as cells treated with *O. hexasperma* extract and infected or not with SIV (p ranging from 0.0131 to 0.0334).

**Figure 7.**
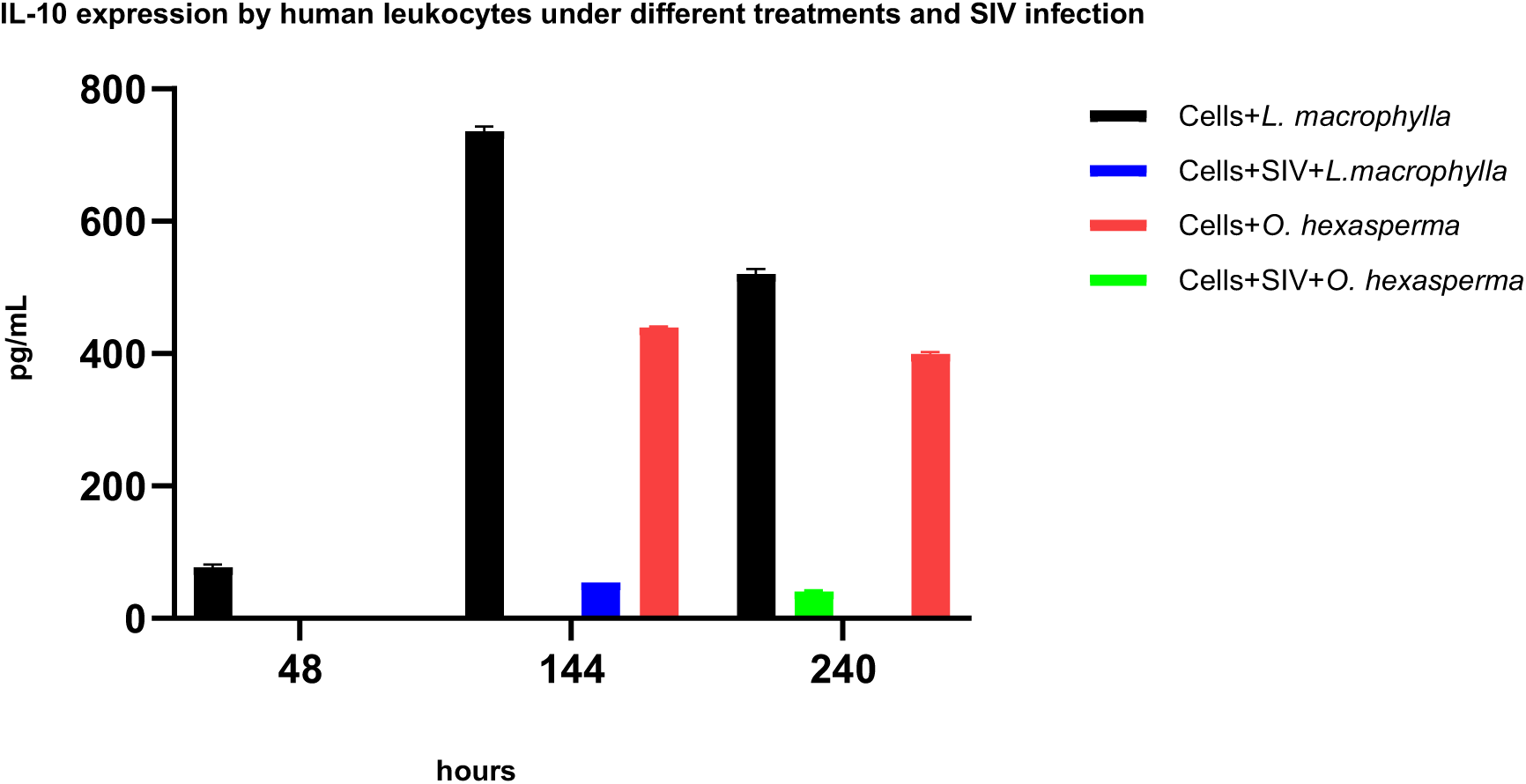
Interleukin 10 expression in established human leukocyte cell line treated with *L. macrophylla* and *O. hexasperma* and SIV infected, and in controls of cells submitted to extracts treatment and no SIV infection.

After 48 hs incubation period, cells treated with *L. macrophylla* extract produced significatively more IFN-γ (Fig. 8) than cells treated with *L. macrophylla* extract and SIV infection (p=0.0028). Other treatments did not show any significative statistical differences (p ranging from 0.0530 to 0.7585) among them (*O. hexasperma* extract treatment versus *O. hexasperma* extract treatment and SIV infection; *O. hexasperma* extract treatment versus *L. macrophylla* extract treatment; *O. hexasperma* extract treatment versus *L. macrophylla* extract treatment and SIV infection; *O. hexasperma* extract treatment and SIV infection versus *L. macrophylla* extract treatment; *O. hexasperma* extract treatment and SIV infection versus *L. macrophylla* extract treatment and SIV infection). At the next time interval of 144 hs, cells treated with *O. hexasperma* extract and SIV infected expressed significatively more IFN-γ than cells treated with *L. macrophylla* extract infected or not with SIV (p ranging from 0.0073 to 0.0370) as also in cells treated with *O. hexasperma* extract and cells treated with *L. macrophylla* extract (p ranging from 0.0177 to 0.0252). Other conditions did not show any statistical differences (p ranging from 0.0557 to 0.7666) among treatments (*O. hexasperma* extract treatment versus *L. macrophylla* extract treatment and SIV infection; *L. macrophylla* extract treatment versus *L. macrophylla* extract treatment and SIV infection). At the last 240 hs incubation period, cells treated with *O. hexasperma* extract expressed significatively more IFN-γ than cells treated with *O. hexasperma* extract and infected with SIV, as also cells treated with *L. macrophylla* extract and SIV infected (p ranging from 0.0159 to 0.0475). Cells treated with *L. macrophylla* extract produced significatively more IFN-γ than cells treated with *L. macrophylla* extract and SIV infected (p=0.0338). Other treatments comparison (*O. hexasperma* extract treatment and SIV infection versus *L. macrophylla* extract treatment and SIV infection; *O. hexasperma* extract treatment and SIV infection versus *L. macrophylla* extract treatment; *O. hexasperma* extract treatment versus *L. macrophylla* extract treatment) did not show any statistical significative difference (p ranging from 0.0887 to 0.2624).

**Figure 8.**
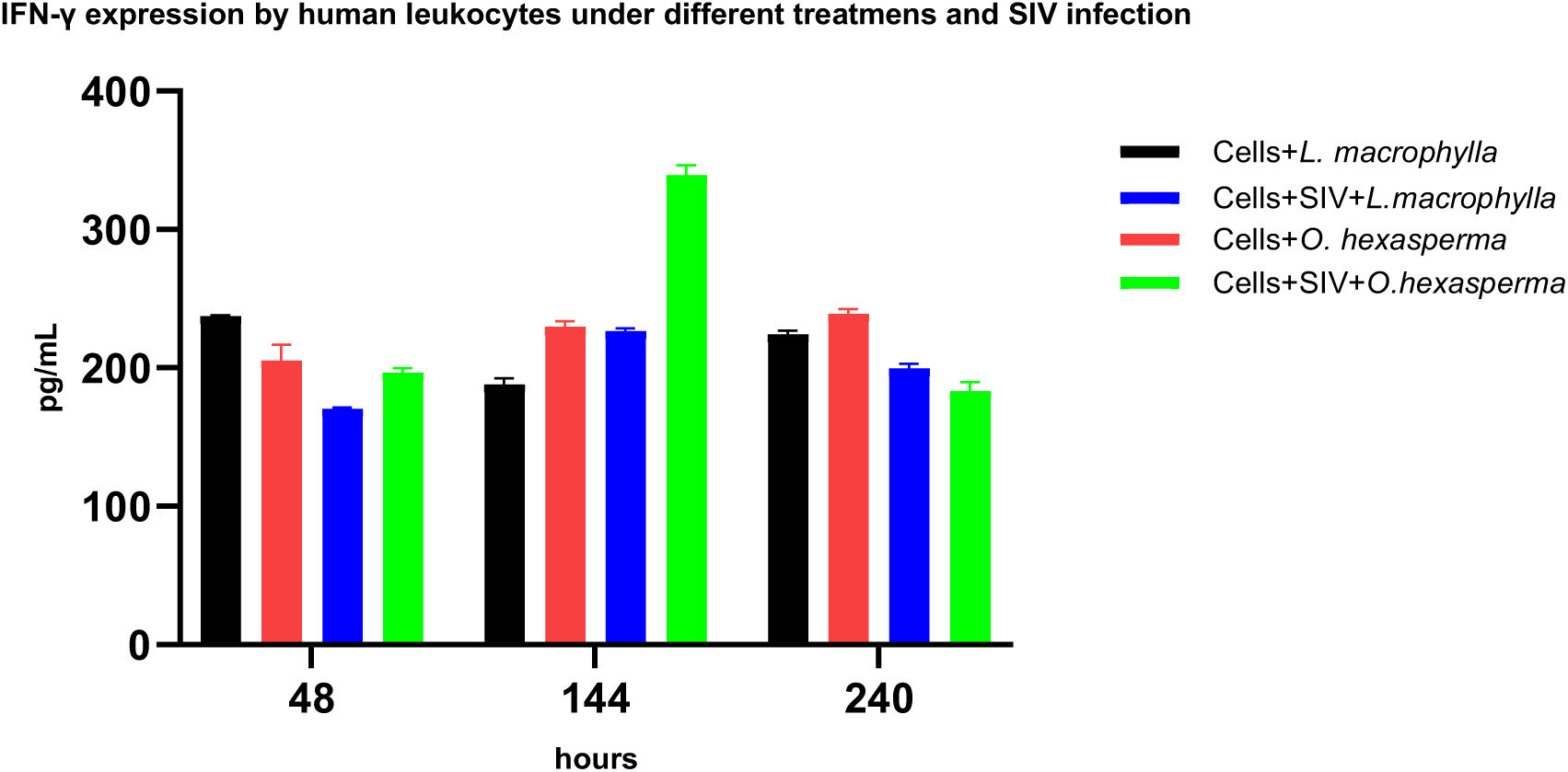
Interferon gamma expression in established human leukocyte cell line treated with *L. macrophylla* and *O. hexasperma* and SIV infected, and in controls of cells submitted to extracts treatment and no SIV infection.

## DISCUSSION

Lentivirus infection, in both humans and nonhuman primates, increases the expression of inflammatory cytokines, prominently IL-8, concomitantly to the elevated SIV/HIV viral proteins synthesis during the virus cycle replication [27], thus so far IL-8 high expression during all time intervals, until 240 hs (Fig. 6), indicates persistent SIVmac_251_ replication and chronicity of infection (Fig. 3) as it happens to occur naturally among lentivirus host infected subjects, mainly among non-treated ones, as also in vivo experiments reported by other authors. Important to emphasize that HIV/SIV *vpr* gene product is responsible for the elevated expression of IL-6, IL-8 and IL-10 among other cytokines in primate leukocyte cells as utilized in our experiments [29]. Besides, IL-8 and IL-6 expression are positively correlated to clinical symptoms in HIV/AIDS patients and disease aggravation [30,31]. Interesting results we obtained, corroborating Dwivedi et al. study [32] of a cohort of HIV reservoir patients, in which HIV unspliced RNA was found to be significantly associated to IL-10 signaling, plus other factors as TLR4/microbial translocation and IL-1/NRLP3 inflammasome. In our experiments, IL-10 was poorly expressed in SIV infected cells (Fig. 7), previously treated or not with plant extracts, but exuberantly secreted by uninfected cells, treated or not with plant extracts, leading to the interpretation that in our in vitro system, HIV unspliced RNA is rare or absent, therefore if lower IL-10 production by EHLC, treated with *L. macrophylla* and *O. hexaspema* extracts, and challenged with SIV infection triggers lentivirus RNA splicing, the compounds of these plants could be potentially associated candidates to antiretroviral therapy stimulating HIV replication in latently infected cells in drug resistant patients [33].

Taking somewhat as a reference to our results, despite in vivo, the clinical research work of Ka’e et al. [34] among HIV-1 infected adolescents in Cameroon, under antiretroviral therapy, quantified a high expression of pro-inflammatory cytokines, IFN-γ and IL-6, besides anti-inflammatory cytokines IL-4 and IL-10. Summing up to other parameters, including also from uninfected subjects, it was stated that the inflammatory cytokines, IFN-γ and IL-12, the anti-inflammatory cytokines, IL-4 and IL-10, and inflammation related cytokines, IL-6 and IL-1β, were markers of successful viral suppression when comparing to viral failure. In our study, we observed that both *O. hexasperma* and *L. macrophylla* plant extracts, infected or not with SIVmac_251_, induced IFN-γ production, being highly and significatively expressed (p=0.0073) by EHLC, treated with *O. hexasperma* and infected with SIV, in the time interval of 144 hs, comparing to other treatments. Concerning IL-6, both plant extracts, under different treatments induced its expression, with significatively higher expression by uninfected EHLC treated with *O. hexasperma* extract than to other treatments with and without SIV infection. Overall, SIV infected leukocytes produced more IL-6 when treated with *O. hexasperma* extract than with *L. macrophylla* extract. Related to the anti-inflammatory cytokines, SIV infected leukocytes treated with *O. hexasperma* extract expressed significatively more IL-4 than other treatments in all time intervals analyzed (p ranging from 0.0208 to 0.0040). Secretion of IL-10 by EHLC was restricted to cells treated with *L. macrophylla* extract in all time intervals (p ranging from 0.0156 to 0.0131). SIV infected leukocytes even treated with plant extracts expressed very low levels of IL-10. Just in the time intervals of 144 hs and 240 hs, IL-10 expression was detected in cells treated with *O. hexasperma* extract. In the initial time of experiment, cells treated with plant extract did produce more cytokines than solely SIV infected cells (data not shown), except for IL-6 that there was not any significative statistical difference. Therefore, our in vitro results are in agreement to the data obtained with the clinical study with HIV-1 infected adolescents.

Whereas betulinic acid has been described for several *Lycania* species, such as *L. tomentosa*, *L. heteromorpha*, *L. carii*, *L. pyrifolia*, *L. cruegeriana*, and *L. licaniaeflora* [35–39], niruflavone was so far only isolated from *L. arianeae* [40].

Many authors found that derivatives of betulinic acid inhibited in vitro the cytopathic effects of HIV cell syncytia formation, suggesting that the anti-HIV activity occurs at virus glycoprotein cell receptor interaction [41–43], lately corroborated by Aiken and Chen [44], which reported the in vitro fusion inhibition of HIV glycoprotein to host cell receptor as also hampering virion maturation, both mechanisms exerted by betulinic acid, which we have isolated from *L. macrophylla*. Niruriflavone is known for its antioxidant activity and has not been associated to antiretroviral activity [45]. The other compounds we obtained, (-)gallocatechine and 4’-O-methyl-epigallocatechin, were shown to inhibit the HIV-1 reverse transcriptase [46–48].

Despite, in our work, the crude extract of *O. hexasperma* was not fractionated, there are many reports of their isolated compounds [23,24]. Among them, the bioflavonoids, agathisflavone, epicathechin, 6-C-glycopyranosyl-luteolin and amentoflavone, even from different plants, exhibited in vitro anti-HIV activity by distinct mechanisms [49,50]. In addition, luteolin and apigenin inhibit HIV enzymes, as the protease, RNase H and integrase [50,51].

## CONCLUSIONS

Our experiments preliminarily demonstrate the mitotic, immune-modulating and SIV suppressive properties of Amazonian medicinal extracts of the plants *L. macrophylla* and *O. hexasperma*. For the most part, *O. hexasperma* extract had the best mitotic activity, while *L. macrophylla* extract was the most protective against SIV cytopathicity. Concerning the cytokines secretion in SIV infected EHLC, *O. hexasperma* extract was generally the most effective to induce their expression, except when referring to IL-8, that there was no difference at all, in the role of both plants’ extracts. The IL-10 was poorly stimulated by both plants’ extract in infected cells. The SIV suppressive activity by the plants’ compounds can result from direct or indirect mechanisms, either by interfering in the lentivirus infection cycle and/or by the action of pro-inflammatory cytokines. Further, restriction factors expressed by leukocytes antagonize the virus pathogenesis. Therefore, both the cell and the lentivirus model to screen antiviral and immune-modulating cytokines are an accessory tool to prospect future therapeutics, not only for retroviruses but also for other viral agents.

## AUTHOR CONTRIBUTIONS

SMSN, JPRAB, BGT, CMC, BSLD, RMB, JFOS and ECGM conducted the experiments and identified the medicinal plants. SSC and IK supervised, conceptualized and obtained grants. IK wrote and edited the manuscript.

## ACKNOWLEDGEMENTS

We are indebted to ImmunoTools GmbH company for the donation of ELISA kits. This research work was funded by FAP DF (Grant number under the process 00193-00000210/2019-60) and University of Brasilia (Grant number under the process 23106.103923/2021-72).

## REFERENCES

1. Atanasov AG, Waltenberger B, Pferschy-Wenzig EM, Linder T, Wawrosch C, Uhrin P, Temml V, Wang L, Schwaiger S, Heiss EH, Rollinger JM, Schuster D, Breuss JM, Bochkov V, Mihovilovic MD, Kopp B, Bauer R, Dirsch VM, Stuppner H. Discovery and resupply of pharmacologically active plant-derived natural products: A review. Biotechnol Adv. 2015;33(8):1582–1614. doi: 10.1016/j.biotechadv.2015.08.001.

2. Arribas P, Andújar C, Bidartondo MI, Bohmann K, Coissac É, Creer S, deWaard JR, Elbrecht V, Ficetola GF, Goberna M, Kennedy S, Krehenwinkel H, Leese F, Novotny V, Ronquist F, Yu DW, Zinger L, Creedy TJ, Meramveliotakis E, Noguerales V, Overcast I, Morlon H, Vogler AP, Papadopoulou A, Emerson BC. Connecting high-throughput biodiversity inventories: Opportunities for a site-based genomic framework for global integration and synthesis. Mol Ecol. 2021;30(5):1120–1135. doi:10.1111/mec.15797.

3. Duarte EA, Novella IS, Weaver SC, Domingo E, Wain-Hobson S, Clarke DK, Moya A, Elena SF, de la Torre JC, Holland JJ. RNA virus quasispecies: significance for viral disease and epidemiology. Infect Agents Dis. 1994;3(4):201–14.

4. Ben-Shabat S, Yarmolinsky L, Porat D, Dahan A. Antiviral effect of phytochemicals from medicinal plants: Applications and drug delivery strategies. Drug Deliv Transl Res. 2020;10(2):354–367. doi: 10.1007/s13346-019-00691-6.

5. Carr A, Mackie NE, Paredes R, Ruxrungtham K. HIV drug resistance in the era of contemporary antiretroviral therapy: A clinical perspective. Antivir Ther. 2023;28(5):13596535231201162. doi: 10.1177/13596535231201162.

6. Menéndez-Arias L, Delgado R. Update and latest advances in antiretroviral therapy. Trends Pharmacol Sci. 2022;43(1):16–29. doi: 10.1016/j.tips.2021.10.004.

7. Berrut G, de Decker L. Immunosénescence: une revue [Immunosenescence: a review]. Geriatr Psychol Neuropsychiatr Vieil. 2015;13 Suppl 2:7–14. French. doi: 10.1684/pnv.2015.0548.

8. Sun L, Wang X, Saredy J, Yuan Z, Yang X, Wang H. Innate-adaptive immunity interplay and redox regulation in immune response. Redox Biol. 2020;37:101759. doi: 10.1016/j.redox.2020.101759.

9. Wautier JL, Wautier MP. Pro- and Anti-Inflammatory Prostaglandins and Cytokines in Humans: *A Mini Review*. Int J Mol Sci. 2023;24(11):9647. doi: 10.3390/ijms24119647.

10. Chang Y, Durante KM. Why consumers have everything but happiness: An evolutionary mismatch perspective. Curr Opin Psychol. 2022;46:101347. doi: 10.1016/j.copsyc.2022.101347.

11. Chang J, Jiang T, Shan X, Zhang M, Li Y, Qi X, Bian Y, Zhao L. Pro-inflammatory cytokines in stress-induced depression: Novel insights into mechanisms and promising therapeutic strategies. Prog Neuropsychopharmacol Biol Psychiatry. 2024;131:110931. doi: 10.1016/j.pnpbp.2023.110931.

12. Zindel J, Kubes P. DAMPs, PAMPs, and LAMPs in Immunity and Sterile Inflammation. Annu Rev Pathol. 2020;15:493–518. doi: 10.1146/annurev-pathmechdis-012419-032847.

13. Çiçek SS, Pfeifer Barbosa AL, Wenzel-Storjohann A, Segovia JFO, Bezerra RM, Sönnichsen F, Zidorn C, Kanzaki I, Tasdemir D. Chemical and Biological Evaluation of Amazonian Medicinal Plant *Vouacapoua americana* Aubl. Plants (Basel). 2022;12(1):99. doi: 10.3390/plants12010099.

14. Pavi CP, Prá ID, Cadamuro RD, Kanzaki I, Lacerda JWF, Sandjo LP, Bezerra RM, Segovia JFO, Fongaro G, Silva IT. Amazonian medicinal plants efficiently inactivate Herpes and Chikungunya viruses. Biomed Pharmacother. 2023;167:115476. doi: 10.1016/j.biopha.2023.115476.

15. Çiçek SS, Galarza Pérez M, Wenzel-Storjohann A, Bezerra RM, Segovia JFO, Girreser U, Kanzaki I, Tasdemir D. Antimicrobial Prenylated Isoflavones from the Leaves of the Amazonian Medicinal Plant *Vatairea guianensis* Aubl. J Nat Prod. 2022;85(4):927–935. doi: 10.1021/acs.jnatprod.1c01035.

16. Oliveira AA, Segovia JF, Sousa VY, Mata EC, Gonçalves MC, Bezerra RM, Junior PO, Kanzaki LI. Antimicrobial activity of amazonian medicinal plants. Springerplus. 2013;2:371. doi: 10.1186/2193-1801-2-371.

17. da Mata ECG, Gonçalves MCA, Segovia JFO, Bezerra RM, Carvalho JCT, Kanzaki LIB. Antiretroviral activity of Amazonian plants. Retrovirology. 2011;8(Suppl 2):P87. doi: 10.1186/1742-4690-8-S2-P87.

18. Sothers CA, Prance GT. *Hymenopus macrophyllus* (Benth). Kew Bull. 2016; 71(4)-58:20.

19. Queiróz, JAL, Mochiuti S, Machado SA. Características silviculturais e potencial de uso da espécie arbórea Licania macrophylla Benth (anoerá/anauerá). Macapá: Embrapa Amapá [Forestry characteristics and potencial utilization of the arboreal species Licania macrophylla Benth]. 2005. Available at: <https://www.infoteca.cnptia.embrapa.br/infoteca/handle/doc/341761>

20. Medeiros FA de, Medeiros AAN, Tavares JF, Barbosa Filho JM, Lima E de O, Silva MS da. Licanol, um novo flavanol, e outros constituintes de Licania macrophylla Benth [Licanol, a new flavanol, and Other constituents from the Licania macrophylla Benth]. Quím Nova [Internet]. 2012;35(6):1179–83. Available at<10.1590/S0100-40422012000600021>

21. Correia AF, Segovia JF, Gonçalves MC, de Oliveira VL, Silveira D, Carvalho JC, Kanzaki LI. Amazonian plant crude extract screening for activity against multidrug-resistant bacteria. Eur Rev Med Pharmacol Sci. 2008;12(6):369–80.

22. Lemos, J. R. (2020) Morfoanatomia de plantas do semiárido (Morphoanatomy of semiarid plants) – 1. ed. – São Paulo: Blucher Open Access, 84 p. il. Available at: < https://pdf.blucher.com.br/openaccess/9786555060485/completo.pdf>.

23. Daniel JF de S, Carvalho MG de, Cardoso R da S, Agra M de F, Eberlin M N. (2005). Others flavonoids from Ouratea hexasperma (Ochnaceae). J. Braz. Chem. Soc. 2005; 16(3b), 634–638. doi.org/10.1590/S0103-50532005000400022

24. Fidelis QC, Faraone I, Russo D, Aragão Catunda-Jr FE, Vignola L, de Carvalho MG, de Tommasi N, Milella L. Chemical and Biological insights of *Ouratea hexasperma* (A. St.-Hil.) Baill.: a source of bioactive compounds with multifunctional properties. Nat Prod Res. 2019;33(10):1500–1503. doi: 10.1080/14786419.2017.1419227.

25. Grynberg NF, Carvalho MG, Velandia JR, Oliveira MC, Moreira IC, Braz-Filho R, Echevarria A. DNA topoisomerase inhibitors: biflavonoids from Ouratea species. Braz J Med Biol Res. 2002;35(7):819–22. doi: 10.1590/s0100-879x2002000700009.

26. da Mata ECG, Kanzaki LIB. A Simple Method to Immortalize Human Leukocytes and its Potential Applications. Microbiol Infect Dis. 2019; 3(2): 1–3.

27. da Mata ECG, Ombredane A, Joanitti GA, Kanzaki LIB, Schwartz EF. Antiretroviral and cytotoxic activities of Tityus obscurus synthetic peptide. Arch Pharm (Weinheim). 2020;353(11):e2000151. doi: 10.1002/ardp.202000151.

28. Gazdar AF, Carney DN, Bunn PA, Russell EK, Jaffe ES, Schechter GP, Guccion JG. Mitogen requirements for the in vitro propagation of cutaneous T-cell lymphomas. Blood. 1980;55(3):409–17.

29. Roux P, Alfieri C, Hrimech M, Cohen EA, Tanner JE. Activation of transcription factors NF-kappaB and NF-IL-6 by human immunodeficiency virus type 1 protein R (Vpr) induces interleukin-8 expression. J Virol. 2000;74(10):4658–65. doi: 10.1128/jvi.74.10.4658-4665.2000.

30. Ellwanger JH, Valverde-Villegas JM, Kaminski VL, de Medeiros RM, Almeida SEM, Santos BR, de Melo MG, Hackenhaar FS, Chies JAB. Increased IL-8 levels in HIV-infected individuals who initiated ART with CD4^+^ T cell counts <350 cells/mm^3^ - A potential hallmark of chronic inflammation. Microbes Infect. 2020;22(9):474–480. doi: 10.1016/j.micinf.2020.05.019.

31. Cummings MJ, Bakamutumaho B, Price A, Owor N, Kayiwa J, Namulondo J, Byaruhanga T, Jain K, Postler TS, Muwanga M, Nsereko C, Nayiga I, Kyebambe S, Che X, Sameroff S, Tokarz R, Shah SS, Larsen MH, Lipkin WI, Lutwama JJ, O’Donnell MR. HIV infection drives pro-inflammatory immunothrombotic pathway activation and organ dysfunction among adults with sepsis in Uganda. AIDS. 2023;37(2):233–245. doi: 10.1097/QAD.0000000000003410.

32. Dwivedi AK, Gornalusse GG, Siegel DA, Barbehenn A, Thanh C, Hoh R, Hobbs KS, Pan T, Gibson EA, Martin J, Hecht F, Pilcher C, Milush J, Busch MP, Stone M, Huang ML, Reppetti J, Vo PM, Levy CN, Roychoudhury P, Jerome KR, Hladik F, Henrich TJ, Deeks SG, Lee SA. A cohort-based study of host gene expression: tumor suppressor and innate immune/inflammatory pathways associated with the HIV reservoir size. PLoS Pathog. 2023;19(11):e1011114. doi: 10.1371/journal.ppat.1011114.

33. Schonhofer C, Yi J, Sciorillo A, Andrae-Marobela K, Cochrane A, Harris M, Brumme ZL, Brockman MA, Mounzer K, Hart C, Gyampoh K, Yuan Z, Montaner LJ, Tietjen I. Flavonoid-based inhibition of cyclin-dependent kinase 9 without concomitant inhibition of histone deacetylases durably reinforces HIV latency. Biochem Pharmacol. 2021;186:114462. doi: 10.1016/j.bcp.2021.114462.

34. Ka’e AC, Nanfack AJ, Ambada G, Santoro MM, Takou D, Semengue ENJ, Nka AD, Bala MLM, Endougou ON, Elong E, Beloumou G, Djupsa S, Gouissi DH, Fainguem N, Tchouaket MCT, Sosso SM, Kesseng D, Ndongo FA, Sonela N, Kamta ACL, Tchidjou HK, Ndomgue T, Ndiang STM, Nlend AEN, Nkenfou CN, Montesano C, Halle-Ekane GE, Cappelli G, Tiemessen CT, Colizzi V, Ceccherini-Silberstein F, Perno CF, Fokam J. Inflammatory profile of vertically HIV-1 infected adolescents receiving ART in Cameroon: a contribution toward optimal pediatric HIV control strategies. Front Immunol. 2023;14:1239877. doi: 10.3389/fimmu.2023.1239877.

35. Fernandes J, Castilho RO, da Costa MR, Wagner-Souza K, Coelho Kaplan MA, Gattass CR. Pentacyclic triterpenes from Chrysobalanaceae species: cytotoxicity on multidrug resistant and sensitive leukemia cell lines. Cancer Lett. 2003;190(2):165–9. doi: 10.1016/s0304-3835(02)00593-1.

36. Braca A, Morelli I, Mendez J, Battinelli L, Braghiroli L, Mazzanti G. Antimicrobial triterpenoids from Licania heteromorpha. Planta Med. 2000 Dec;66(8):768–9. doi: 10.1055/s-2000-9601.

37. Bilia, A.R., Mendez J, Morelli I (1996a) Phytochemical investigations of Licania genus. Flavonoids and triterpenoids from Licania carii. Pharm Acta Helv 71,191–197. 10.1016/0031-6865(96)00010-6

38. Estrada O, Contreras W, Acha G, Lucena E, Venturini W, Cardozo A, Alvarado-Castillo C. Chemical constituents from Licania cruegeriana and their cardiovascular and antiplatelet effects. Molecules. 2014;19(12):21215–25. doi: 10.3390/molecules191221215.

39. Braca A, Sortino C, Mendez J, Morelli I. Triterpenes from Licania licaniaeflora. Fitoterapia. 2001;72(5):585–7. doi: 10.1016/s0367-326x(00)00321-x.

40. Carvalho MG de, Costa PM da. Outros constituintes isolados de Licania arianeae (Chrysobalanaceae) [Other constituents isolated from Licania arianeae (Chrysobalanaceae)]. Rev. Bras. Farmacogn. 2009; 19(1b), 290–293. 10.1590/S0102-695X2009000200018

41. Fujioka T, Kashiwada Y, Kilkuskie RE, Cosentino LM, Ballas LM, Jiang JB, Janzen WP, Chen IS, Lee KH. Anti-AIDS agents, 11. Betulinic acid and platanic acid as anti-HIV principles from Syzigium claviflorum, and the anti-HIV activity of structurally related triterpenoids. J Nat Prod. 1994;57(2):243–7. doi: 10.1021/np50104a008.

42. Mayaux JF, Bousseau A, Pauwels R, Huet T, Hénin Y, Dereu N, Evers M, Soler F, Poujade C, De Clercq E, et al. Triterpene derivatives that block entry of human immunodeficiency virus type 1 into cells. Proc Natl Acad Sci U S A. 1994;91(9):3564–8. doi: 10.1073/pnas.91.9.3564.

43. Hashimoto F, Kashiwada Y, Cosentino LM, Chen CH, Garrett PE, Lee KH. Anti-AIDS agents--XXVII. Synthesis and anti-HIV activity of betulinic acid and dihydrobetulinic acid derivatives. Bioorg Med Chem. 1997;5(12):2133–43. doi: 10.1016/s0968-0896(97)00158-2.

44. Aiken C, Chen CH. Betulinic acid derivatives as HIV-1 antivirals. Trends Mol Med. 2005;11(1):31–6. doi: 10.1016/j.molmed.2004.11.001.

45. Than NN, Fotso S, Poeggeler B, Hardeland R, Laatsch H. (Niruriflavone, a New Antioxidant Flavone Sulfonic Acid from Phyllanthus niruri. ChemInform. 2006; 37(25). doi: 10.1002/chin.200625196

46. Moore PS, Pizza C. Observations on the inhibition of HIV-1 reverse transcriptase by catechins. Biochem J. 1992;288 (Pt 3)(Pt 3):717–9. doi: 10.1042/bj2880717.

47. Hussein G, Miyashiro H, Nakamura N, Hattori M, Kawahata T, Otake T, Kakiuchi N, Shimotohno K. Inhibitory effects of Sudanese plant extracts on HIV-1 replication and HIV-1 protease. Phytother Res. 1999;13(1):31–6. doi: 10.1002/(SICI)1099-1573(199902)13:1<31::AID-PTR381>3.0.CO;2-C.

48. Lin YM, Anderson H, Flavin MT, Pai YH, Mata-Greenwood E, Pengsuparp T, Pezzuto JM, Schinazi RF, Hughes SH, Chen FC. In vitro anti-HIV activity of biflavonoids isolated from Rhus succedanea and Garcinia multiflora. J Nat Prod. 1997;60(9):884–8. doi: 10.1021/np9700275.

49. Nath S, Bachani M, Harshavardhana D, Steiner JP. Catechins protect neurons against mitochondrial toxins and HIV proteins via activation of the BDNF pathway. J Neurovirol. 2012;18(6):445–55. doi: 10.1007/s13365-012-0122-1.

50. Mehla R, Bivalkar-Mehla S, Chauhan A. A flavonoid, luteolin, cripples HIV-1 by abrogation of tat function. PLoS One. 2011;6(11):e27915. doi: 10.1371/journal.pone.0027915.

51. Tang TT, Li SM, Pan BW, Xiao JW, Pang YX, Xie SX, Zhou Y, Yang J, Wei Y. Identification of Flavonoids from *Scutellaria barbata* D. Don as Inhibitors of HIV-1 and Cathepsin L Proteases and Their Structure-Activity Relationships. Molecules. 2023;28(11):4476. doi: 10.3390/molecules28114476.

